# A neural circuit for competing approach and avoidance underlying prey capture

**DOI:** 10.1101/2020.08.24.265181

**Authors:** Daniel Rossier, Violetta La Franca, Taddeo Salemi, Cornelius T. Gross

## Abstract

Predators must frequently balance competing approach and avoidance behaviors elicited by a moving and potentially dangerous prey. Several brain circuits supporting predation have recently been localized. However, the mechanisms by which these circuits balance the conflict between approach and avoidance responses remain unknown. Laboratory mice initially show alternating approach and avoidance responses toward cockroaches, a natural prey, but with repeated exposure become avid hunters. Here we used *in vivo* neural activity recording and cell-type specific manipulations in hunting mice to identify neurons in the lateral hypothalamus and periaqueductal grey that encode and control predatory approach and avoidance. We found a subset of GABAergic neurons in lateral hypothalamus that specifically encoded hunting behaviors and whose stimulation triggered predation, but not feeding. This population projects to the periaqueductal grey and stimulation of these projections promoted predation. Neurons in periaqueductal grey encoded both approach and avoidance behaviors, but only initially when the mouse showed high levels of fear of the prey. Our findings allow us to propose that GABAergic neurons in lateral hypothalamus facilitate predation in part by suppressing defensive responses to prey encoded in the periaqueductal grey. Our findings reveal a neural circuit mechanism for controlling the balance between conflicting approach and avoidance behaviors elicited by the same stimulus.

## Introduction

The ability to seek and capture prey adeptly is a conserved behavior essential to the survival of numerous animal species. However, attacking prey brings risks for the predator and efficient hunting requires a skillful balance between responding to threatening and appetitive prey cues. The neural circuits involved in this balance are unknown. Rodents naturally hunt and consume a variety of insects and hunting behavior has been studied in the laboratory using rats given cockroaches or mice (1) and more recently in laboratory mice given crickets (2-5). In early studies, immediate early gene mapping during hunting identified the recruitment of lateral hypothalamus (LHA) (1, 6, 7) and periaqueductal grey (PAG) (7, 8) and electrical stimulation of either LHA (9-12) or PAG (13, 14) could initiate avid predatory attacks in rats and cats suggesting the presence of neuronal cell populations that promote hunting in both these brain structures.

LHA has been historically regarded as a central node in the neural system controlling seeking behaviors, including feeding and predatory hunting (14-16). Recent circuit neuroscience work has confirmed this role by finding that stimulation of *Vgat*+ GABAergic neurons in LHA can trigger predation as well as feeding (5, 17-20). However, the role of these neurons in predation may be indirect, as any manipulation that increases food-seeking is expected to promote motivation to hunt as well. Moreover, some studies in which LHA GABAergic neurons were optogenetically stimulated did not observe the full repertoire of feeding behaviors, but instead found increased chewing activity, digging, or general motor activity (21-23), suggesting that the link between LHA-driven feeding and hunting may be more complex. One possible confound in these studies is their use of two different Cre driver lines (*Vgat*:Cre vs. *Gad2*::Cre) that have been shown to target distinct LHA GABAergic neurons (23). Thus, it remains unclear to what extent different populations of LHA GABAergic neurons specifically encode and control predatory behaviors.

A link between LHA and PAG in predation was made by a recent study showing that activation of LHA to PAG projections can promote predatory hunting in mice (5) and the central importance of the PAG in hunting is further strengthened by several studies identifying a series of PAG afferents arising from different brain structures – including the central nucleus of the amygdala, medial preoptic area, and zona incerta – that promote predation (3, 4, 24, 25). However, the precise functional role of the PAG in hunting remains unclear. For example, it was found that inhibition of glutamatergic neurons in the lateral and ventrolateral PAG (l/vlPAG) by GABAergic inputs from the central nucleus of the amygdala promotes predatory behavior (3) suggesting that PAG neurons might function to suppress rather than promote predation. At the same time, other studies have shown that activation of glutamatergic neurons in l/vlPAG induces defensive behaviors (26) a finding that is consistent with an extensive literature linking PAG to both innate and learned defensive behavior (27-29). One explanation for these observations is that PAG has an antagonistic role in hunting by promoting avoidance of the prey early in the encounter when defensive responses to prey are maximal. Suppression of these defensive responses by GABAergic inputs would thus facilitate unimpeded predation.

Here we identify *Gad2*+ neurons in LHA as those specifically recruited during predatory chasing and attack and both sufficient and necessary to drive hunting, but not feeding or social behavior. Activation of LHA *Gad2*+ neuron projections to PAG decreased defensive responses to prey and promoted hunting during the early phases of predation when mice learned to hunt cockroaches. Finally, *in vivo* calcium endoscopy identified a neural population in PAG that encoded risk assessment and flight behaviors elicited by prey. Notably, we failed to detect PAG neurons responsive to predatory pursuit or attack. These results point to a circuit in which activity in LHA *Gad2*+ neurons promotes hunting in part by suppressing avoidance responses encoded in PAG.

## Results

### Mice learn to efficiently hunt cockroaches

To understand the contribution of the lateral hypothalamic seeking system to hunting behavior we studied mice pursuing, capturing, and consuming cockroaches, a natural prey. Exposure of naïve mice to cockroaches elicited repeated attempts to approach and attack the prey interspersed with robust flight and freezing responses (**Figure 1A, Video 1**). Repeated exposure of mice to cockroaches led to a gradual decrease in defensive responses and latency to pursue and capture prey (**Figures 1A-E**) suggesting that either their fear of the prey diminishes or their motivation to hunt is increased with repeated exposure. In trained animals, hunting typically initiated in 43 ± 11 seconds after the introduction of the prey compared to 391 ± 109 seconds in untrained animals. A predatory sequence usually involved bouts of pursuit and attack using mouth and forepaws, followed by attempts to consume the immobilized prey. Residual movement of the immobilized prey during feeding occasionally interrupted the engagement and led to the initiation of new hunting bouts. Importantly, mice fed laboratory chow *ad libitum* showed robust predatory hunting behavior arguing for a strong hunger-independent motivation for mice to pursue and capture prey. The alternation between defensive and predatory behavior toward the cockroach and its evolution over training days allowed us to investigate the recruitment of neural circuits involving the balance between approach and avoidance of prey during hunting.

**Figure 1.**
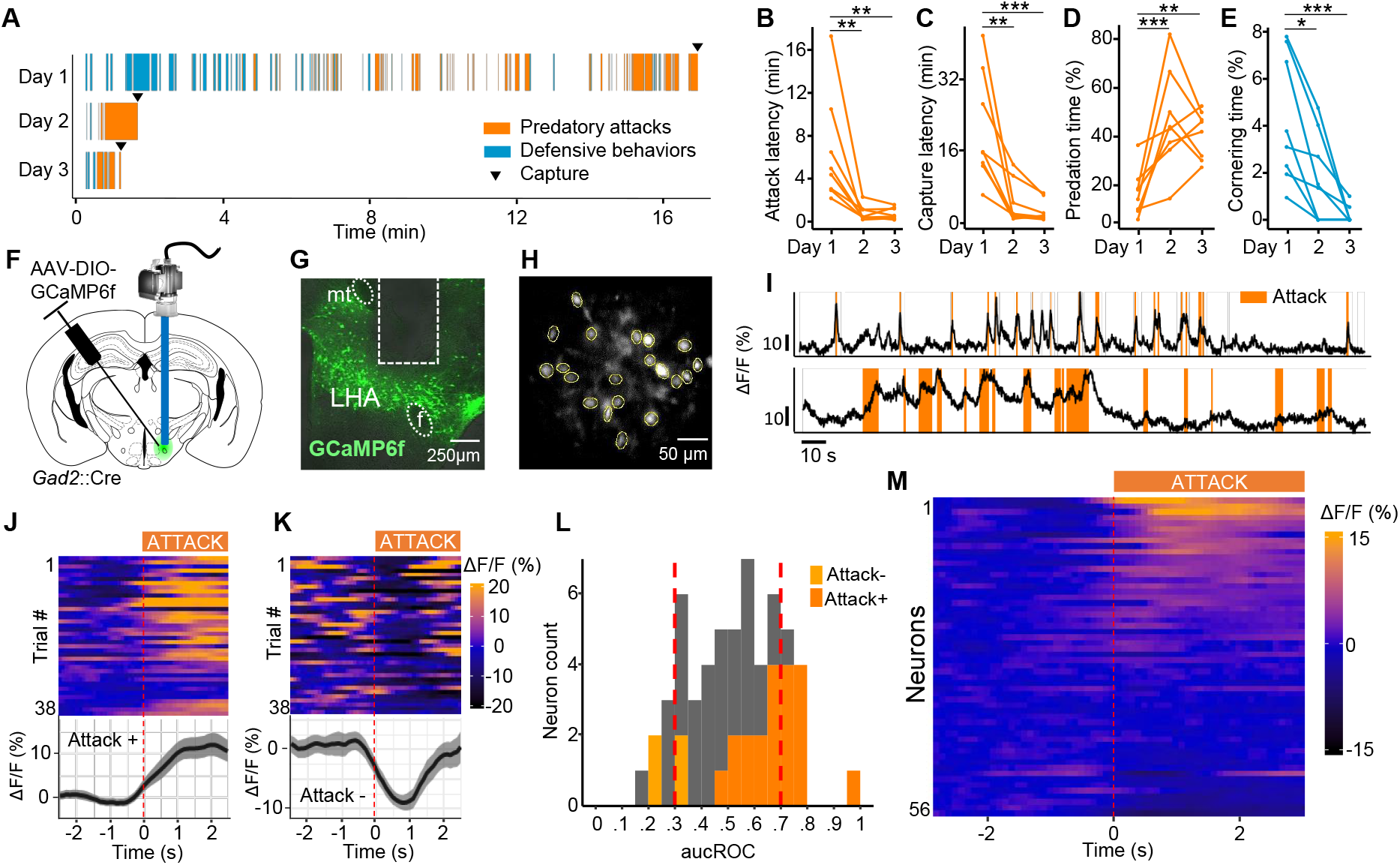
LHA *Gad2*+ neurons are recruited during predatory attack. (**A**) Raster plot illustrating the occurrence of aggressive and defensive behaviors of a representative mouse over three consecutive days of predation training. (**B-E**) Quantification of defensive and predatory behavior over training days (N=8). (**F**) *Gad2*+ transgenic mice were infected with a Cre-dependent GCaMP6f-expressing virus and activity of LHA *Gad2*+ neurons was recorded in living animals using microendoscopy (N=5). (**G**) Representative section showing GRIN lens placement (*mt* mamillary tract, *F* fornix). (**H**) Representative post-processed calcium imaging field of view with ROIs indicated by yellow circles. (**I**) Activity trace (ΔF/F) of two representative cells recruited during predatory attack. (**J**) Response of a cell whose activity was significantly increased following attack onset (Attack+, top: activity heat map where each line corresponds to a single attack event aligned to onset, bottom: average activity trace of 38 attack events aligned to attack onset; ΔF/F ± SEM). (**K**) Response of a cell whose activity was significantly decreased following attack onset (Attack-, top: activity heat map where each line corresponds to a single attack event, bottom: average activity trace of 38 flight events aligned to attack onset). (**L**) Histogram of area under the receiver operator curve (auROC) values for all cells (56 cells, N = 5). ROI fluorescence per frame is used to classify attack vs. non-attack periods. Vertical lines in histogram indicate high (0.7) and low (0.3) cut-off (Attack+: dark orange, Attack-: light orange). (**M**) Peri-attack activity heatmap for all cells aligned to attack onset. Cells are ranked by activity increase at attack onset (N = 5, Bonferroni *post hoc* test: *P < 0.05, **P < 0.01, ***P < 0.001).

### Lateral hypothalamus neurons encode hunting

Because the previous report that *Gad2*+ neurons were linked to physical activity, but not feeding (23) we investigated whether this subclass of GABAergic neuron is recruited during predation. For long-term monitoring of neural activity, mice were surgically implanted with a GRIN lens cannula adapted to fit a head-mounted miniature fluorescent microscope following viral delivery of the genetically encoded calcium sensor GCaMP6f (AAV-*EF1a*::DIO-GCaMP6f) in the LHA (**Figure 1FH** and **S1A, Video 2**). We expressed the calcium sensor in *Gad2*+ neurons using a Cre-dependent virus delivered to *Gad2*::Cre driver mice. Following 5 to 9 days of training, surgically treated mice showed robust and repeated hunting episodes with very few defensive responses to prey. A significant number of recorded neurons showed activity patterns that correlated with predatory attack of prey across hunting episodes (**Figure 1I**). To identify neurons with significant hunting-correlated activity, calcium activity signals surrounding the initiation of predatory attacks were superimposed across episodes and statistically assessed (N = 5 mice, 12-38 episodes). A subset of neurons showed a significant positive (20/56, 36%) or negative (4/56, 7%) correlation with attack initiation (**Figure 1JK, M**). The significance of these correlations was confirmed by a bootstrap analysis where we replaced the attack events by randomized events (**Figure S1B**). Subsets of neurons showed a global decrease (6/56, 11%) or increase (10/56, 17%, corresponding to <0.3 and >0.7 auROC score, respectively) in activity that was greater during episodes of hunting relative to other behaviors as assessed by analysis of receiver operator characteristic (ROC) curves (**Figure 1L**). As expected, neurons showing increased activity to the onset of predation also tended to show increased responses during hunting episodes in the ROC analyses (**Figure 1L**). These observations confirm that predatory chasing and attack behaviors recruit neural activity in LHA *Gad2*+ neurons.

### *Lateral hypothalamus* Gad2*+ neurons are necessary and sufficient for hunting*

To determine whether activity in *Gad2*+ neurons in LHA is necessary and sufficient for hunting behavior we used optogenetic activation and pharmacogenetic inhibition to selectively increase and decrease neuronal activity in the presence of a cockroach. To test whether neuronal activity in LHA *Gad2*+ neurons is sufficient to increase hunting behavior we optogenetically stimulated the LHA of untrained *Gad2*::Cre mice that had been bilaterally infected with AAV-*Ef1a*::DIO-ChR2-EYFP virus and exposed them to a cockroach (**Figure 2A-C** and **S1A**). Light stimulation in ChR2 expressing animals presented for the first time with a cockroach did not affect the initial latency to investigate the prey when compared to control animals (**Figure 2D**). However, light stimulation was associated with a significant decrease in latency to attack the prey in ChR2 expressing vs. control animals (**Figure 2E**) and the total time spent attacking the prey was significantly increased in ChR2 animals during light stimulation epochs or when compared to controls not expressing ChR2 (**Figure 2F** and **Video 3**). Light stimulation in ChR2 animals was also associated with significantly less defensive behavior in which the animal remained in the corner of the cage facing the prey when compared to control mice (**Figure 2G**).

**Figure 2.**
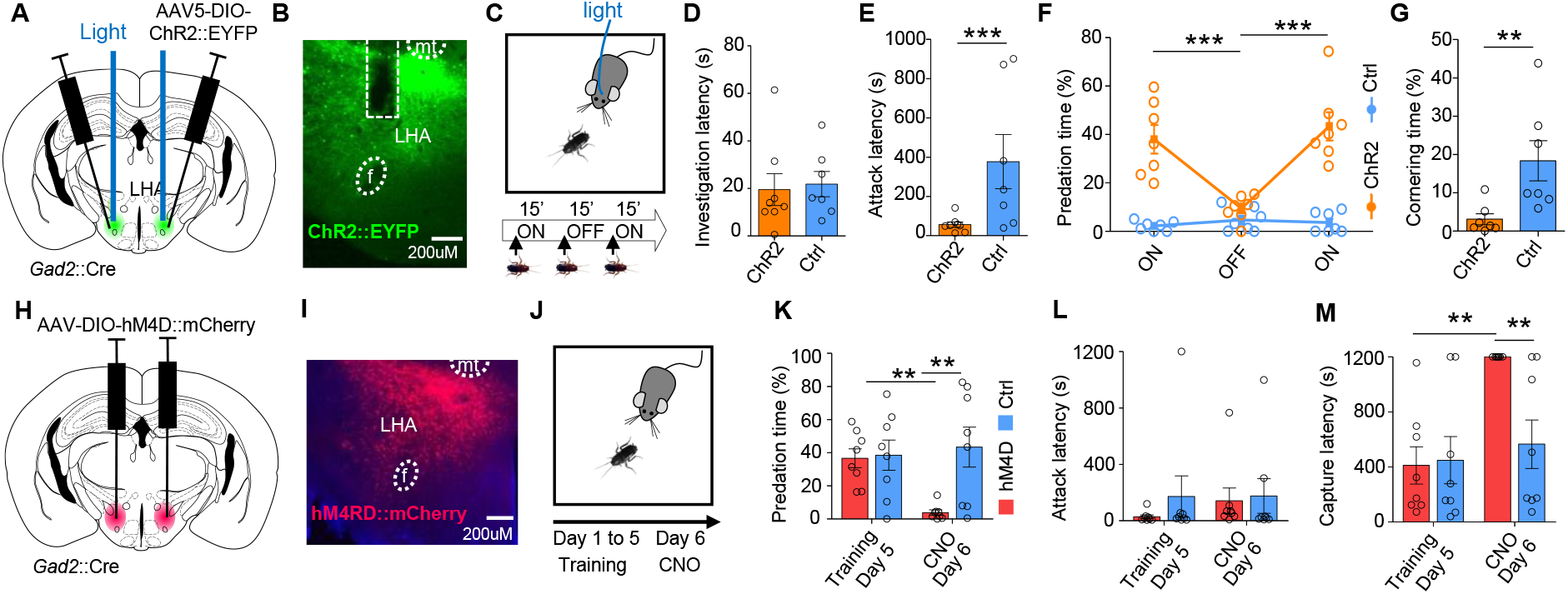
LHA *Gad2*+ neurons promote predation. (**A**) *Gad2*::Cre transgenic mice were infected with a Cre-dependent ChR2-expressing or control virus bilaterally in LHA and optic fibers were bilaterally implanted in the dorsal part of the LHA. (**B**) Representative section showing ChR2 reporter expression and fiber placement in LHA. (**C**) Mice naïve to prey were exposed to a cockroach during two epochs of light stimulation (ON, 15 min) separated by one epoch with no stimulation (OFF, 15 min). (**D**) Latency to begin investigation of prey (t-test P = 0.47). (**E**) Latency to attack prey (t-test P< 0.0001). (**F**) Time spent attacking the prey (ChR2: N = 7, F_(2,22)_ = 13.6, P = 0.0001). (**G**) Time spent cornering during first stimulation epoch, (ChR2: N = 7, t-test, P = 0.0065; unless otherwise indicated, ChR2: N = 8, Ctrl: N = 7, mean ± SEM, repeated-measures ANOVA – epoch x group interaction). (**H**) *Gad2*+ transgenic mice were infected with a Cre-dependent hM4D-expressing or control virus bilaterally in LHA. (**I**) Representative section showing hM4D reporter expression in LHA. (**J**) Animals underwent predation training over five days and were tested on the sixth day following treatment with CNO. (**K**) Time spent attacking on days 5 and 6 (F_(2,14)_ = 11.46, P = 0.0044). (**L**) Latency to attack on days 5 and 6 (F_(2,14)_ = 0.27, P = 0.58). **M)** Latency to capture prey on days 5 and 6 (F_(2,14)_ = 9.83, P = 0.0073; hM4D: N = 8, Ctrl: N = 8, mean ± SEM, repeated measure ANOVA – treatment x group interaction, Bonferroni *post hoc* test *P < 0.05, **P < 0.01, ***P < 0.001, *mt* mamillary tract, *F* fornix).

Finally, we noted that ChR2 animals receiving light stimulation often did not consume the prey, but released it after capture and continued with predatory pursuit, consistent with the idea that LHA *Gad2*+ neurons promote the preparatory rather than consummatory phases of hunting.

To test whether neural activity in LHA *Gad2*+ neurons is necessary to promote hunting *Gad2*::Cre mice were bilaterally infected in the LHA with AAV-*hSyn*::DIO-hM4D-mCherry or control virus (**Figure 2HI** and **S2B**) and animals were trained to hunt efficiently by daily exposure to cockroaches over 5 days. On the sixth day animals were treated with clozapine-N-oxide (CNO) one hour before testing (**Figure 2J**). Although CNO treatment was not associated with a significant change in latency to attack prey (**Figure 2L**), CNO treatment significantly decreased time spent attacking (**Figure 2K**) and increased latency to capture (**Figure 2M**) prey in hM4D expressing animals compared to controls. Together these findings argue for a causal role of LHA *Gad2*+ neurons in promoting hunting, but potentially not feeding.

### Lateral hypothalamus *Gad2*+ neurons do not promote feeding or aggression

To directly examine whether LHA *Gad2*+ neurons are involved in feeding behavior we optogenetically activated this population when animals were given a choice between a chamber where food was available and another where it was not (**Figure 3A**). Light stimulation of ChR2 expressing animals did not elicit increased food consumption or time spent in a food containing chamber when compared to controls (**Figure 3BC**). Nevertheless, we noted that stimulation of ChR2 expressing animals triggered avid biting of food pellets and both Petri dishes (**Figure 3D** and **S2C, Video 4**). These findings confirm that *Gad2*+ neurons do not promote feeding *per se*, but that they can trigger hunting-like capture behavior even towards non-prey objects.

**Figure 3.**
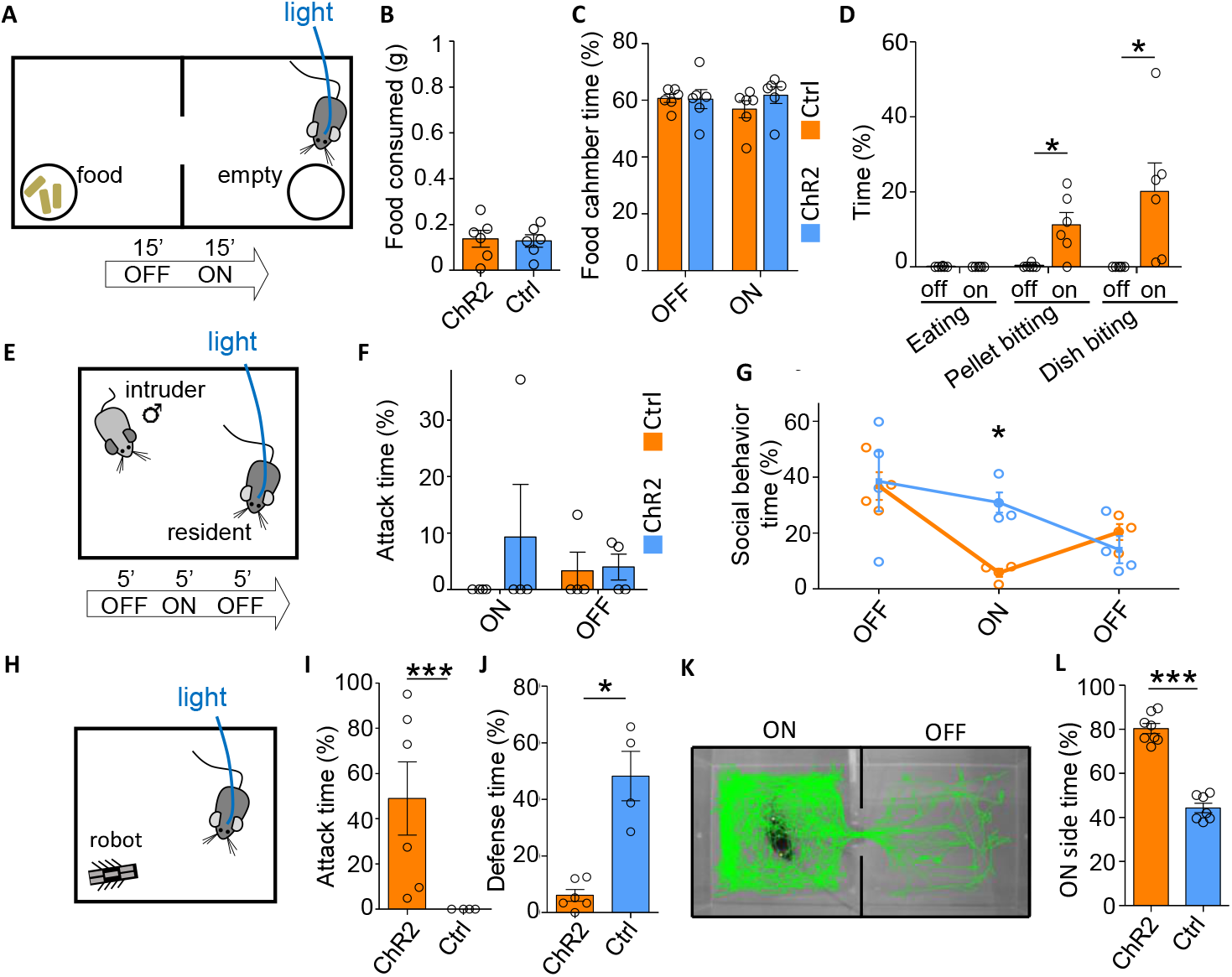
LHA *Gad2*+ neurons do not promote feeding or aggression. (**A**) *Gad2*::Cre mice infected with a Cre-dependent ChR2-expressing or control virus in LHA and implanted with optical fibers were tested in a two compartment apparatus with food available on one side and an empty petri dish on the other side. Following a period without stimulation (OFF, 15 min) animals were stimulated with light (ON, 15 min). (**B**) Food consumed (weight_BEFORE_ - weight_AFTER_, t-test P = 0.84). (**C**) Time spent on food side (F_(2,8)_ = 2.17, P = 0.17). (**D**) Time spent eating or biting food pellet or petri dish (F_(3,36)_ = 5.59, P = 0.0077; ChR2: N = 6, Ctrl: N = 6, mean ± SEM, repeated measure ANOVA - epoch x group). (**E**) To test social aggression a subordinate mouse was introduced into the home cage of the experimental animal. (**F**) Time spent attacking the intruder (F_(2,6)_ = 0.92, P = 0.37). **G)** Percentage of time spent performing social behavior (F_(3,12)_ = 4.37, P = 0.037; ChR2: N = 4, Ctrl: N = 4, mean ± SEM, repeated measure ANOVA - epoch x group). **H)** Real-time place preference (RTPP) assay was use to examine the reinforcing properties of LHA *Gad2*+ neuron stimulation. Representative path tracking in (left) stimulated and (right) non-stimulated compartments for a ChR2-expressing mouse. (**I**) Time spent on the stimulation side of the RTPP apparatus (t-test P < 0.0001; ChR2: N = 8, Ctrl: N = 7, mean ± SEM). (**J**) Predation of a prey-like moving object was examined by introducing a small mechanical robot into the cage. (**K**) Time spent attacking the robot (Mann-Whitney test, P = 0.009). (**L**) Time spent performing defensive behaviors in the presence of the robot (P = 0.0005; ChR2: N = 6, Ctrl: N = 4, mean ± SEM, Bonferroni *post hoc* test: *P < 0.05, **P < 0.01, ***P < 0.001)

Next, we investigated whether stimulation of LHA *Gad2*+ neurons might increase attack against conspecifics. Mice were subjected to the resident-intruder test of social aggression in which a subordinate BALB/c male is introduced into the cage of a singly housed experimental animal (**Figure 3E**). Light stimulation of ChR2 expressing resident mice was not associated with an increase in attacks toward the intruder (**Figure 3F**), we instead observed a decrease in social activity (**Figure 3G**), suggesting that this population does not regulate aggression towards conspecifics. However, we did observe that stimulated mice occasionally chased and bit the tail of intruders, a behavior we did not see in stimulated control animals (**Video 5**).

These findings suggest that stimulation of LHA *Gad2*+ neurons is able to elicit predation-like behaviors (e.g. chasing, biting) towards inanimate objects. To test this hypothesis we examined whether ChR2 stimulation of LHA *Gad2*+ neurons could drive predatory attack against a prey-like robot (**Figure 3J**). Light stimulation of ChR2 expressed in LHA *Gad2*+ neurons was associated with pursuit and attack of the robot, a behavior not observed in control animals (**Figure 3K**). Importantly, control animals showed a much higher level of avoidance of the artificial prey than of a natural prey, reinforcing the conclusion that stimulation of LHA *Gad2*+ neurons is able to overcome defensive behaviors elicited by the prey to promote pursuit (**Figure 3L**).

Finally, we tested whether optogenetic stimulation of LHA *Gad2*+ neurons had rewarding properties in the real-time place preference test (RTPP). Light stimulation of mice expressing ChR2 in LHA *Gad2*+ neurons in one side of a two chamber shuttle box was associated with a significant preference for the stimulated side when compared to control mice (**Figure 3HI**). These findings suggest that LHA *Gad2*+ neurons promote reinforcing behavior.

### Projections from LHA to PAG reduce prey avoidance

We hypothesized that PAG might serve primarily to promote prey avoidance and that inhibitory projections from LHA to PAG could promote hunting by suppressing defensive responses to the prey during the initial learning phase. Consistent with this hypothesis, we confirmed that LHA *Gad2*+ neurons project to the l/vlPAG (**Figure S3A**). To test whether these projections have a functional role in the suppression of defensive responses to prey we bilaterally expressed ChR2 in LHA *Gad2*+ neurons (AAV-*Ef1a*::DIO-ChR2-EYFP) and stimulated their projections in the l/vlPAG (**Figure 4AB** and **S3B**). In untrained ChR2 animals, but not in control mice, light stimulation was associated with a significant reduction in latency to attack **(Figure 4C)**. However, this effect was not associated with a decreased latency to capture compared to controls, suggesting that only part of the predatory behavioral sequence was affected **(Figure 4D)**. Consistent with a decreased latency to initiate predation, total time spent attacking the cockroach was significantly increased in ChR2 compared to control animals. **(Figure 4E)**. Moreover, defensive responses to the cockroach as measured by cornering behavior were significantly reduced in ChR2 mice compared to controls (**Figure 4F**). A similar decrease in defensive behaviors was seen toward the mechanical robot. Notably, in some cases stimulated animals proceeded to attack the robot, something that was only rarely observed in unstimulated or control animals (**Figure 4G**). These findings support a role for *Gad2*+ LHA-PAG projections in promoting predation and suggest that this occurs at least in part via a reduction in defensive response to prey.

**Figure 4.**
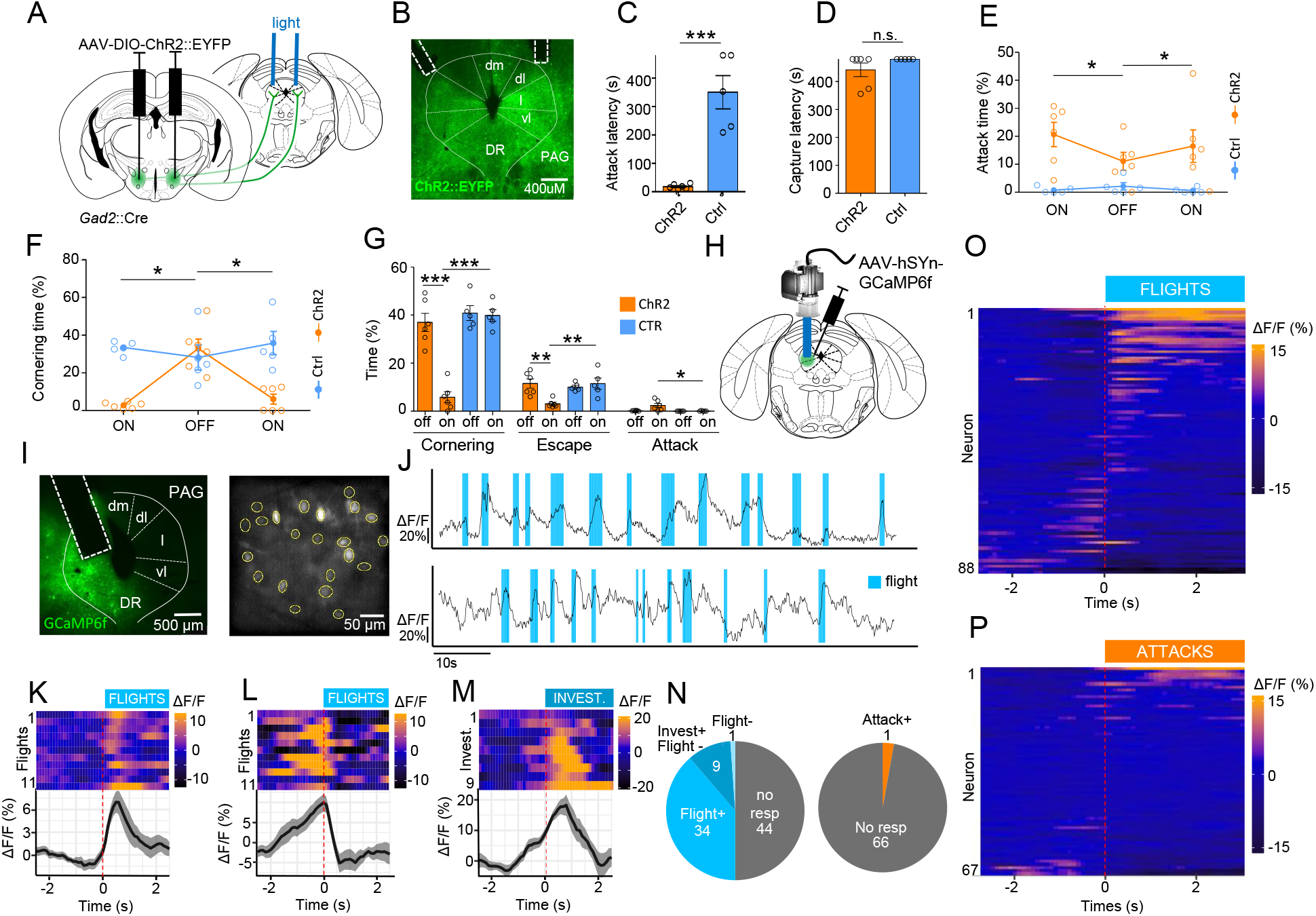
LHA to PAG projections inhibit defensive response to prey. (**A**) *Gad2*::Cre mice infected with a Cre-dependent ChR2-expressing or control virus in LHA and implanted with optical fibers over lPAG were tested for predatory hunting. On the first day of testing animals were subjected to a period without stimulation (OFF, 8 min) followed by light stimulated (ON, 8 min). (**B**) Representative section showing ChR2 reporter expression and fiber placement (*dm*: dorsomedial, *dl*: dorosolateral, *l*: lateral, *vl*: ventrolateral, *DR*: dorsal raphe). (**C**) Latency to attack prey (t-test, P = 0.0001). (**D**) Latency to capture prey (Mann-Whitney test, P = 0.56). (**E**) Time spent attacking prey (F_(3,18)_ = 4.06, P = 0.035). (**F**) Time spent cornering (F_(3,18)_ = 15.6, P = 0.0002; ChR2: N = 6, Ctrl: N = 5, mean ± SEM, repeated measure ANOVA - epoch x group). In a separate group of animals LHA to PAG projection activation was performed in mice presented a prey-like robot. (**G**) Time spent (left) cornering, (center) escaping and (right) attacking in the presence of a mechanical robot. (**H**) Mice were infected with a GCaMP6f-expressing virus and activity of lPAG neurons was recorded in living animals using a microendoscopy (N = 5). (**I**) Representative section with (left) GRIN lens placement and (right) post-processed field of view. (**J**) Activity trace (ΔF/F) of two representative neurons recruited during flight. (**K**) Response of a neuron that significantly increased its activity at flight onset (Flight+). Each line of the heat map (top) corresponds to a single attack event aligned to attack onset. Traces of neuron activity averaged across 11 flight events (bottom) aligned to attack onset (ΔF/F, mean ± SEM). Response of a neuron that (**L**) decreased activity at flight onset (Flight-), or (**M**) increased activity at investigation onset (Assessment+). (top) Each line corresponds to a single attack event and (bottom) average responses across 38 flight events. (**N**) Distribution of neurons significantly increasing (Flight+, Assessment+) or decreasing (Flight-, Assessment-) activity at flight and investigation onset. (**O**) Distribution of neurons significantly increasing (Attack+) and decreasing (Attack-) activity at attack onset recorded in animals following successful training to hunt. Heatmap of average peri-flight neuron activity aligned to flight onset for all neurons recorded in (**P**) naïve (N = 5) or (**Q**) trained (N = 4) mice. (Bonferroni *post hoc* test: *P < 0.05, **P < 0.01, ***P < 0.001).

### Periaqueductal grey neurons encode prey avoidance

To test the hypothesis that PAG neurons might primarily encode defensive response during predation we performed *in vivo* calcium microendoscopy in the lateral PAG (lPAG) of mice exposed repeatedly to a cockroach. GCaMP6f (*AAV-hSyn*::GCaMP6f) was expressed in l/vlPAG and animals were surgically implanted with a microendoscope above lPAG and exposed over several day to a cockroach (**Figure 4HI** and **S3C**). On the first day of training when mice exhibited frequent flights from the prey a significant number of neurons showed activity patterns that were visibly correlated with avoidance behavior **(Figure 4J)**. To identify neurons in PAG with significant behavior-correlated activity calcium activity signals surrounding the initiation of flight episodes were superimposed and statistically assessed. A large fraction of neurons (39%, 34/88) showed a significant increase in activity time-locked to the onset of flight (Flight+; **Figure 4KNO**). A smaller proportion of neurons (11%, 10/88) showed a significant decrease in activity at the on-set of flight (Flight-; **Figure 4LNO**). The significance of these correlations was confirmed by a bootstrap analysis where we replaced the flight events by randomized events (**Figure S3D**). Flights were often preceded by risk assessment behaviors oriented toward the prey, including stretched approach, sniffing, and rearing. Notably, most Flight-cells showed an increase in activity during this risk assessment period, increasing their activity up to the moment of flight onset and then returning to baseline thereafter (**Figure 4L**). Because these units showed significant time-locked activity to the onset of risk assessment behaviors we labeled them as assessment cells (Assessment+; **Figure 4M**).

To understand whether PAG activity was modulated by the level of fear toward the prey we examined the activity of neurons in animals trained to efficiently hunt prey and showing no, or only very infrequent avoidance behaviors (5-9 days of training). Under these conditions mice showed prey pursuit and attack that were not associated with risk assessment or followed by flight. Consistent with the absence of flight behaviors we failed to detect Flight+ cells during or following prey attack, although we did observed one unit whose activity significantly increasing at attack onset (**Figure 4N,P**). Notably, we failed to detect any units whose activity was correlated with approach to prey. These observations suggest that the Assessment+ cell activity seen during approach to prey in untrained animals is not linked to prey approach or predation per se, but rather may encode prey-associated features that are attenuated during repeated exposure or training. These data are consistent with the hypothesis that neurons in lPAG encode defensive rather than predatory behavior.

## Discussion

Our data show that neural activity in a genetically defined subset of GABAergic neurons (*Gad2*+) in the LHA is necessary and sufficient to promote hunting behavior, including both prey pursuit and attack. Importantly, LHA *Gad2*+ neuron stimulation did not promote the consumption of stationary food suggesting that this population specifically drives prey pursuit (**Figure 3BD**). These data contrast with earlier studies which found that LHA GABAergic neurons stimulation could trigger feeding (17, 19, 20). This discrepancy may in part be explained by the observation that the *Gad2*::Cre driver line is expressed in partially non-overlapping neural populations in LHA compared to the *Vgat*::Cre line used in these studies (23). We note that earlier studies had reported that stimulation of *Gad2*+ neurons did not eliciting feeding, but rather a general increase in physical activity (23). In light of our data, we interpret this increased activity as a latent predatory impulse observed in the absence of prey. Alternatively, the discrepancy may be linked to the hedonic value of prey that may arise from a combination of chemosensory and movement cues essential to trigger predation (2, 24, 25). In this case, *Gad2*+ neurons would be selectively involved in increasing the drive to seek prey because of its rewarding nature, while not affecting food seeking because it depends primarily on hunger. However, we don’t favor this interpretation because stimulation of *Gad2*+ neurons elicited attack also against non-prey objects (**Figure 3D**). It remains to be determined whether *Gad2*+ neuron activity might be more responsive to highly appetitive foods or under food deprivation, conditions which were not explored in our study.

Earlier work has shown that LHA GABAergic neurons can promote a variety of approach behaviors in addition to hunting and feeding, including social investigation and aggression and unifying theories of LHA function point to a more general role in supporting the vigor of positively reinforced seeking behaviors (15, 30). However, stimulation of *Gad2*+ neurons in our study did not increase social aggression in the resident-intruder assay, but rather appeared to increase physical activity that interfered with social interaction (**Figure 3G**). Our observation that the resident animal occasionally chased and bit the tail of the intruder during LHA *Gad2*+ stimulation may reflect the promotion of predation-like behaviors toward the intruder at the expense of affiliative social interactions. Thus, our finding that *Gad2*+ stimulation promoted attack toward prey and robot, but not toward a social opponent highlights the distinction between social and predatory aggression. This distinction is also consistent with the proposed role for LHA in supporting the appetitive drive involved in the preparatory phase of predation (16). Finally, our finding that stimulation of *Gad2*+ neurons was reinforcing in the RTPP assay (**Figure 3I**) is consistent with previous studies on LHA GABAergic neurons and underlines the reinforcing nature of predation.

An important aim of our work was to investigate how the balance of predation and fear of prey shifts during repeated exposure. Our findings suggest that this balance depends on the cross-inhibition between LHA neurons that promote the motivation to hunt and PAG neurons that promote prey avoidance. Stimulation of LHA neurons during the initial trials could overcome defensive responses to the prey and increase hunting (**Figure 2FG**) and this could be mimicked by stimulation of their projections to PAG (**Figure 4EF**). The role of PAG in prey avoidance was supported by our discovery of two distinct neuronal populations in PAG whose activity was time-locked with prey assessment and flight, respectively (**Figure 4KL**). The existence of neurons encoding both approach and avoidance appears contradictory with a role for PAG in promoting flight. However, similar populations have been observed in dorsal PAG during approach-avoidance behavior to a natural predator (31, 32) and we hypothesize that these populations coordinate defensive responses to prey by promoting risk assessment and triggering escape.

In conclusion, our data is consistent with a model in which inhibitory afferents to PAG directly suppress the firing of neurons that promote risk assessment and flight and thereby allow fearless predation. This model allows us to propose a role for PAG as a brake, rather than as an accelerator for predation. It follows that PAG may function more widely to antagonize seeking behaviors driven by LHA or other upstream structures. Under such a model, a gradual increase in inhibitory inputs to PAG over the course of repeated exposure to prey, including those from LHA, gradually suppresses PAG activity and lowers defensive responses to prey. An alternative hypothesis is that inhibitory afferents to PAG promote predation by suppressing activity in PAG that tonically antagonizes approach behavior. However, although we cannot rule out that low levels of tonic activity escaped detection in our *in vivo* calcium imaging experiments, we do not favor this hypothesis as we did not observe cells in PAG whose activity was suppressed during prey pursuit or attack. Our findings may be relevant to understanding several recent studies examining the role of PAG afferents in predation. Based on our data, we hypothesize that GABAergic inputs from CeA and ZI that promote hunting may do so by blocking defensive behaviors elicited by prey rather than by promoting predation directly. More work aimed at manipulating and recording these circuits in animals during the entire span of predatory learning will be required to test whether our findings generalize across PAG afferent circuits.

## Materials & Methods

### Animals and behavioral apparatus

All experimental procedures involving the use of animals were carried out in accordance with EU Directive 2010/63/EU and under the approval of the EMBL Animal Use Committee and Italian Ministry of Health License 541/2015-PR to C.G. Animals were singly housed in temperature and humidity-controlled cages with *ad libitum* access to food and water under a 12h/12h light-dark cycle. All experiments were carried out in male C57BL/6J mice aged 2-8 months and bred at EMBL. Transgenic mice were heterozygous for *Gad2*::Cre (Gad2^tm2(cre)Zjh^, Taniguchi et al. 2011). All tested animals were singly housed either from the date of surgery or from one week before testing for animals not undergoing surgery and the pharmacogenetic experiments. Subordinate mice in the resident-intruder test were BALB/c males mice aged 7 weeks old and bred at EMBL. Prey were adult female Turkestan cockroaches (*B. Lateralis*, 2-3 cm) purchased from commercial pet supply distributors in Italy and propagated at EMBL.

### Surgical procedures

Mice were anesthetized with 3% isoflurane (Provet) and subsequently head fixed in a stereotaxic frame (Kopf) with body temperature maintained at 36 ± 2 C and anesthesia sustained with 1-2% isoflurane and oxygen. The skull was exposed, cleaned with hydrogen peroxide (0.3% in water), and leveled. Craniotomy was performed with a handheld drill and, when needed, extra holes were drilled to fit 2 or 3 implant-stabilizing miniature screws (RWD). For optogenetic activation experiments 0.2-0.3 μl *AAV5-Ef1a::DIO-* ChR2(E123T/T159C)-EYFP or AAV5-*Ef1a*::DIO-EYFP (UNC Vector Core and Addgene, respectively) virus was injected bilaterally in the LHA via a pulled glass capillary attached to a pneumatic injection system (AP -1.9, ML 0.95, DV 5.5 from Bregma). Optic fibers were implanted bilaterally in the dorsal part of the LHA (0.66 NA, 200 μm, Prizmatix; 230/1250 μm internal/external diameter ceramic ferrule, Thorlabs). Implants were attached to the skull using dental cement (Duralay) and secured with screws fixed to the skull. One of the two fibers was implanted with an angle due to space constraints and to prevent the bilateral lesion of overlying brain structures (LHA: 15° inclination, AP -1.9, ML 1.6, DV 5.5; AP - 1.9, ML 0.95, DV 5.5; PAG: 26° inclination, AP -4.4, ML 1.8, DV 2.7; AP: -4.4, ML 0.5, DV 2.2). For *in vivo* calcium endoscopy 0.2-0.3 *μ*l of AAV5-*hSyn*::DIO-GCaMP6f virus (Penn Vector Core) was infused in the LHA (AP -1.9, ML 0.95, DV 5.5) of *Gad2*::Cre mice or AAV5-*hSyn*::GCaMP6f was infused in the l/vlPAG (AP -4.4, ML 0.6, DV 2.6). Endoscope GRIN lens imaging cannulas (LHA: model E, PAG: model L, type V, Doric Lenses) were implanted in the LHA (AP -1.9, ML 0.95, DV: 5.0) and lPAG (20° inclination, AP -4.4, ML 1.6, DV 2.6) and implants attached to the brain using dental cement (Duralay). Approximately 1 hour before the end of surgery an analgesic was administered (Carprofen 5 mg/kg). Immediately after surgery mice were intraperitoneally injected with 0.4 ml saline and allowed to recover in heated cages. For 3 days post-surgery drinking water was supplemented with paracetamol (10 ml/l). For pharmacogenetic experiments AAV8-*hSyn*::DIO-hM4D-ires-mCherry and AAV8-*hSyn*::DIO-mCherry (UNC Vector Core and Addgene, respectively) virus were infused in the LHA (AP -1.9, ML 0.95, DV 5.5).

### Hunting test

The cage cover holding food and water was removed and replaced with a filter paper lid before mice were tested for predation in their home cage. An adult cockroach was placed in the cage until capture occurred. If prey was captured in less than 4 minutes, the cockroach was replaced. Cockroaches were removed before the mouse could progress to eat the prey, although it was not always possible to prevent consumption. Behaviors were recorded for scoring using cameras placed on the side and/or top of the cage.

### In vivo *calcium microendoscopy*

Expression of GCaMP6f was allowed to proceed for at least 4 weeks following infection. Mice were habituated over three days to the testing cage and the procedure of microscope (Doric Lenses) plugging before testing commenced. Mice in the endoscope experiments were somewhat slower to achieve efficient hunting (5-9 days). During each training day several recording sessions lasting 2-15 minutes were carried out. LED intensity for GCaMP6f stimulation was adapted to each implant (15-60 % max power, 10 Hz). Side and/or top cameras were used to track behavior. For LHA recordings, data was successfully collected from 9 out of 18 animals that underwent surgery. Only animals with >9 distinguishable neurons and histologically confirmed fiber placement were included in the analysis (5/9). For PAG recordings, data was successfully collected from 11 out of 18 animals (5/11 with >9 neurons and correct fiber placement).

### Optogenetic stimulation

Viral expression was allowed to continue for at least 2 weeks (4 weeks for projection activation). Mice were habituated over three days to handling and optic fiber plugging before testing commenced. Optical stimulation was performed with double LED modules (465 nm, Plexbright, Plexon) directly attached to a rotary joint and connected to the implant via a patch cable (1 m, 0.66 NA, Plexbright High Performance, Plexon). Power at the tip of the implanted fiber was verified before surgery with an optical power meter (Thorlabs). Stimulation trains were generated using V2.2 Radiant software (Plexon; LHA: 20Hz, 10ms, 13-17 mW; PAG: 20Hz, 20ms, 13-17 mW). For the hunting test the animal had no prior experience with prey and initiation of stimulation was concomitant with presentation of the cockroach. For technical reasons two ChR2 expressing animals could be analyzed for only the first 15’ stimulation epochs. For the feeding test the mouse was placed in a two compartment plexiglass cage (25 x 50 cm). The mouse was plugged 10 min before being placed in the apparatus containing two Petri dishes, one filled with food pellets and the other empty, placed in each compartment, respectively. For the resident intruder test a male BALB/c intruder (6-7 week old) was placed in the home cage of the tested animal. For the real-time place preference assay animals were plugged 10 min before being placed in a two chamber apparatus. For the robot experiments a Hexbug Nano device was used.

### Pharmacogenetic inhibition

Virus expression was allowed to proceed for three weeks. Testing took place in the home cage following training during which mice were allowed to hunt and consume two cockroaches per day for five consecutive days. One hour before each training session animals received an intraperitoneal saline injection. Behavior was scored on the last training day as baseline. On the testing day all animals received CNO (3 mg/kg, i.p.) one hour before exposure to prey.

### Behavioral data analysis

Behavioral scoring was performed manually using Solomon Coder software at 100 or 200 ms bin frame rate with the experimenter blind to calcium trace or experimental group. Behaviors were defined as – *flight:* rapid and stereotyped running away from prey or robot; *cornering:* immobile in cage corner facing prey (frequently observed following *flight); jump escape:* repeated jumping against cage walls (frequently observed when prey approached cornered mouse); a*ttack:* aggressive behavior towards prey or robot using forepaws or mouth (usually coinciding with horizontal straightened tail); *capture:* prey immobilization and biting; *investigation:* sniffing environment, often while rearing; *social behavior:* anogenital and facial sniffing and back biting (Note: tail biting was not considered social behavior); *eating:* grabbing food using both forepaws and/or mouth chewing; *food pellet biting:* biting without grabbing or chewing.

### Histology

Animals were anesthetized Avertin (2.5%, i.p.) and trans-cardially perfused with PBS followed by 4% PFA (Sigma) in 0.1M PB solution. The brain was then extracted from the skull, post-fixed in 4% PFA for 24 hours at 4°C and sectioned (20-60 μm) using a vibratome (Leica VT 1000s). If necessary, sections were stored in PBS 0.01% sodium azide. For viral transgene expression or implant track position verification sections were mounted on slides and imaged using an epifluorescence or confocal microscope.

### Calcium imaging analysis

Image processing was performed with ImageJ/Fiji software. Videos were batch processed using a customized macro. The background of each frame was calculated using an FFT bandpass filter (lower/higher band: 100/10,000, Fiji) and subtracted from the original images frame by frame. The resulting stack was then aligned with Fiji TurboReg and Neuronal cell body like structures showing variation in fluorescence intensity were manually identified as regions of interest (ROIs, elliptical areas). Several ROIs were designated in areas not containing any visible neurons as negative controls. The average fluorescence intensity of ROIs for each frame were extracted (F). ROIs were not tracked across days. Δ F/F (F-F_0_/F_0_) was calculated for each ROI where F_0_ was the mean fluorescence intensity of the ROI over the entire recording period. A few isolated frames (<1/1000) appeared completely black due to technical issues and ROIs of these frames were interpolated from adjacent frames. ROIs were included in the analysis only if they had at least one transient response, calculated as 3 consecutive frames during the whole recording session being at least twice the standard deviation of the preceding 5 s. To eliminate interference between brief and adjacent behavioral events, and to account for GCaMP6f decay time, events that occurred twice within a 3 s time window were excluded. The peri-behavioral traces were centered on the intensity mean of 5s preceding the behavior onset. Average responses for each neuron were obtained by averaging the normalized intensities across all events and smoothed using a 4-point rolling mean. Statistics to identify cells responding to a given behavior were performed on the average response of at least 9 trials. For ROC analyses, all frames from selected calcium imaging recording sessions were classified as “attack” or “no attack”. A ROC curve was generated for each neuron by plotting the true positive and false positive rates across the distribution and the auROC calculated with 0.3 and 0.7 selected as thresholds for classification.

### Statistical analysis

Prism Graphpad 5 software or custom R scripts were used to generate graphs and perform statistical analyses. Unless otherwise indicated for optogenetic and pharmacogenetic experiments repeated measures ANOVA with Bonferroni post-hoc test as well as t-test were used. For calcium imaging behavioral time locked response adjusted t-test was used. The boot strap analyses were performed by randomly generating, for each recording, a scrambled set of events and measuring the P-value of all the neurons from the recording.

## Acknowledgments

We thank the EMBL Laboratory Animal Facility, Genetic & Viral Engineering Facility (GAVEF), Microscope Facility, and Francesca Zonfrillo, Claudia Valeri, Roberto Voci, Valerio Rossi and Matteo Gaetani for support with animal husbandry and management and Piotr Krzywkowski and Maria Esteban Masferrer for support with calcium imaging and Angelo Raggioli for production of viruses. The work was supported by EMBL (D.R., V.L.F, T.S., C.T.G.), SNF (D.R.), and the European Research Council (ERC) Advanced Grant COREFEAR (C.T.G).

## Author contributions

Behavioral experiments and their analysis were carried out by D.R. with support from V.L.F. Calcium imaging experiments were carried out by D.R. with support from T.S. The project was conceived by D.R. and C.T.G., who both designed the experiments and wrote the manuscript.

---

**Video 1.** Representative defensive and predatory behaviors of mice towards a cockroach.

**Video 2.** Representative defensive and predatory behaviors of *Gad2*::cre mice implanted with microendoscope in LHA.

**Video 3.** Representative predatory behaviors induced by LHA *Gad2*+ neuron stimulation.

**Video 4.** Representative effect of LHA *Gad2*+ neuron stimulation in the feeding test.

**Video 5.** Representative tail biting induced by LHA *Gad2*+ neuron stimulation.

## Supplementary Figure Legends

**Figure S1.**
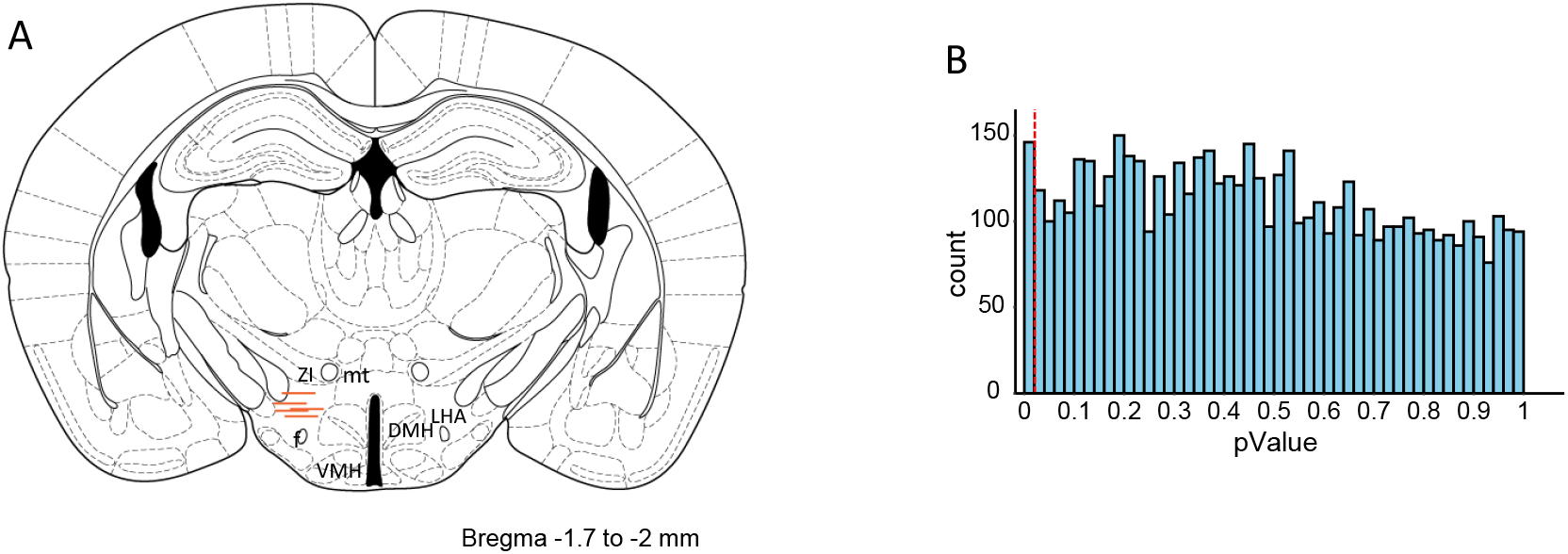
(**A**) Schematic of GRIN lens Placements in LHA. LHA: *lateral hypothalamus*; f: *fornic*; mt: *mamiliray tract*; ZI : *zona incerta*; VMH: *ventromedial hypothalamus*; DMH: *dorsomedial hypothalamus*. (**B**) Histogram of the P-values obtained in a bootstrap test. The positions of the attack events in each recording were scrambled and the P-value of the change in activity at the randomized events calculated for each neuron. The process was repeated 100 times. The 100 P-values of each neuron are reported in the histogram. The vertical red dashed line represents the P = 0.02 cutoff.

**Figure S2.**
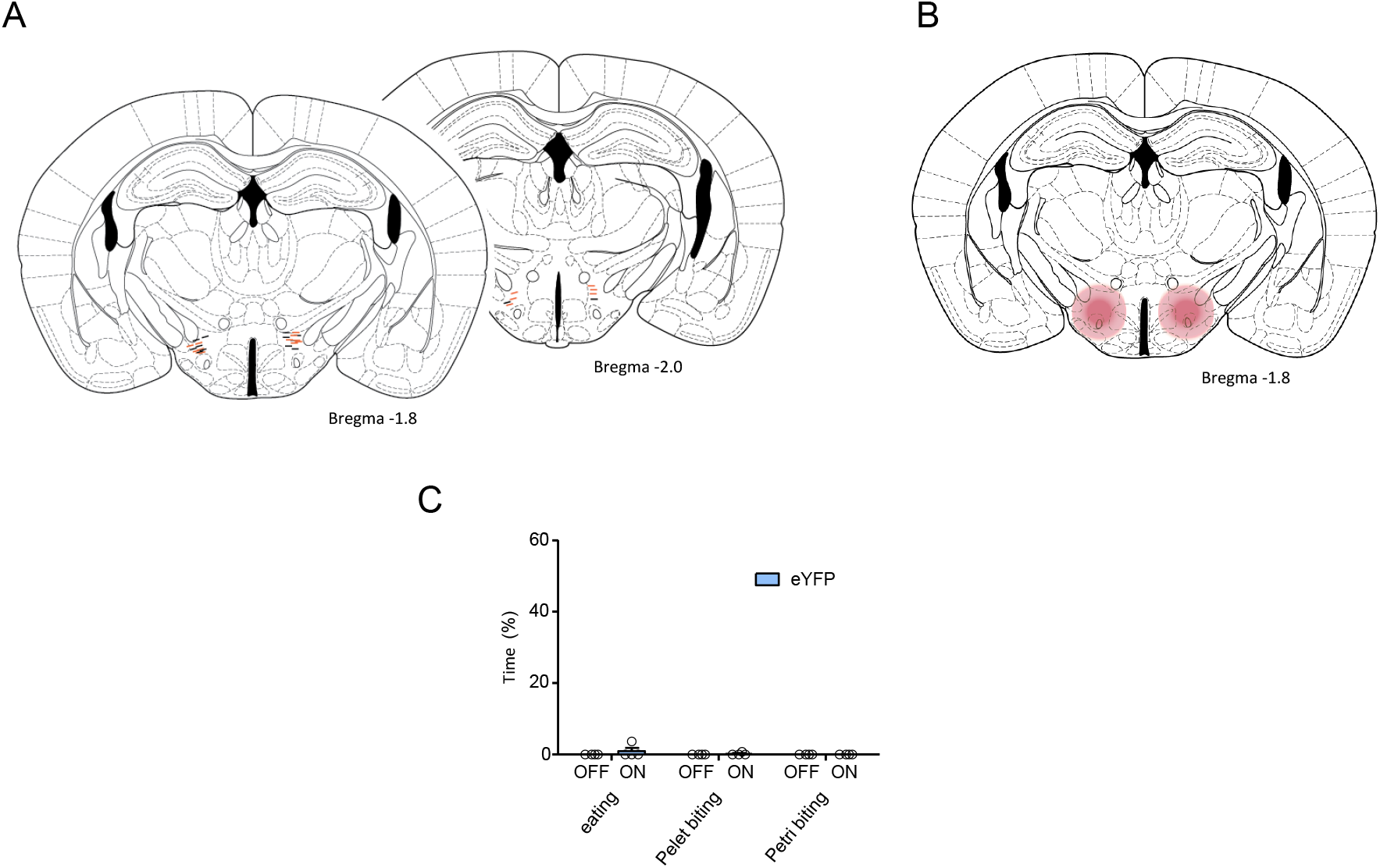
(**A**) Schematic of optic fibers placements in the LHA (Orange: ChR2, Black: Ctrl). (**B**) Schematic of injection area of AAV-*hSyn*::DIO-hM4D-ires-mCherry and AAV-*hSyn*::DIO-mCherry in LHA (dark red: estimated injection area, light red: virus spreading area). (**C**) Percentage of time spent eating (left) and biting the food pellet (center) or petri dish (left) in control animals.

**Figure S3.**
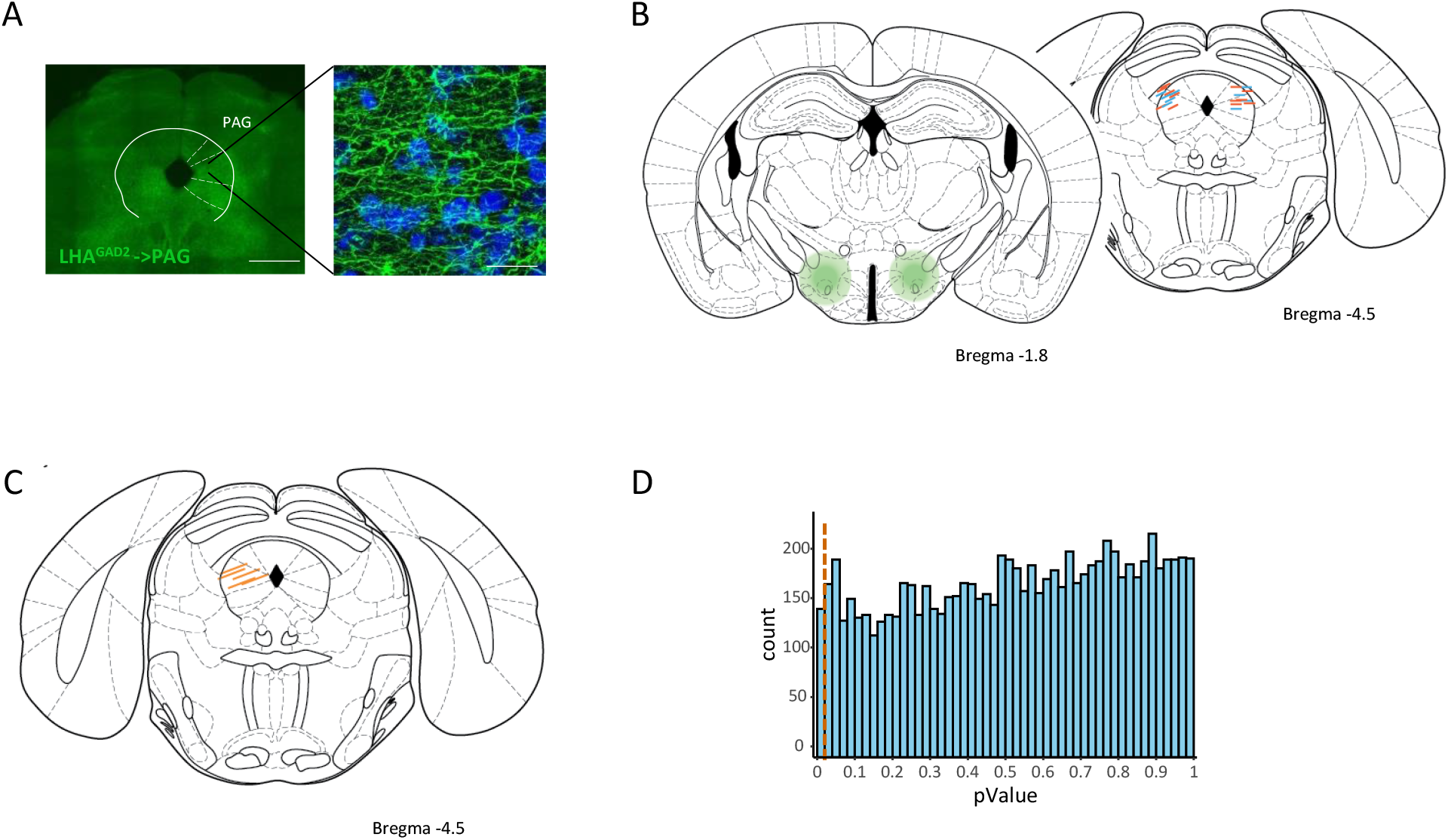
(**A**) PAG section showing ChR2 reporter expression from LHA *Gad2*+ projections. Right magnification of in the lPAG. Blue: DAPI staining. (**B**) Schematic of AAV5-*Ef1a*::DIO-ChR2-EYFP and AAV5-*Ef1a*::DIO-EYFP injection area in the LHA (left). (dark green: estimated injection area, light green: virus spreading area); Optogenetic fibers placement in the PAG (right). (Orange: ChR2, Blue: ctrl). (**C**) Schematic of GRIN lens placements in the PAG. (**D**) Histogram of P-values obtained in a bootstrap test. The positions of the attack events in each recording were scrambled and the P-value of the change in activity at the randomized events calculated for each neuron. The process was repeated 100 times. The 100 P-values of each neuron are reported in the histogram. The vertical red dashed line represents the P = 0.02 cutoff.

